# Development of antibody-dependent cell cytotoxicity function in HIV-1 antibodies

**DOI:** 10.1101/2020.09.24.312066

**Authors:** Laura E. Doepker, Sonja Danon, Elias Harkins, Duncan Ralph, Zak Yaffe, Amrit Dhar, Cassia Wagner, Megan M. Stumpf, Dana Arenz, James A. Williams, Walter Jaoko, Kishor Mandaliya, Kelly K. Lee, Frederick A. Matsen, Julie M. Overbaugh

## Abstract

A prerequisite for the design of an HIV vaccine that elicits protective antibodies is understanding the developmental pathways that result in desirable antibody features. The development of antibodies that mediate antibody-dependent cellular cytotoxicity (ADCC) is particularly relevant because such antibodies have been associated with HIV protection in humans. We reconstructed the developmental pathways of six human HIV-specific ADCC antibodies using longitudinal antibody sequencing data. Most of the inferred naïve antibodies did not mediate detectable ADCC. Gain of antigen binding and ADCC function typically required mutations in complementarity determining regions of one or both chains. Enhancement of ADCC potency often required additional mutations in framework regions. Antigen binding affinity and ADCC activity were correlated, but affinity alone was not sufficient to predict ADCC potency. Thus, elicitation of human ADCC antibodies may require mutations that first enable high affinity antigen recognition, followed by mutations that optimize factors contributing to functional ADCC activity.

## Introduction

A major concept underlying HIV vaccine research is that we can learn from the processes that lead to potent antibody responses in natural infection to inform immunogen design. For this reason, there have been several detailed studies of the evolutionary pathways of potent and broad neutralizing antibodies (bnAbs) to HIV (Bonsignori et al., 2017; Bonsignori et al., 2016; Doria-Rose et al., 2014; Landais et al., 2017; Liao et al., 2013; MacLeod et al., 2016; Simonich et al., 2019; Wu et al., 2015). These studies highlight the critical role of somatic hypermutation (SHM), particularly in complementary determining regions (CDRs) of antibody heavy chains, in driving the breadth and potency of HIV bnAbs (Kwong & Mascola, 2018). One recent study also highlighted that maturation of the antibody light chain can also be critical for breadth (Simonich et al., 2019). Collectively, these studies have provided important insights into the specific mutations that drive maturation of antibody lineages and specificities of bnAbs.

The keen interest in HIV bnAbs stems in part from a plethora of compelling studies that demonstrate the efficacy of bnAbs in protection of macaques from experimental SHIV infection (Pegu, Hessell, Mascola, & Haigwood, 2017). The data are less convincing for sterilizing protection by HIV-specific non-neutralizing antibodies that mediate antibody-dependent cell cytotoxicity (ADCC) in the same experimental animal models (Fouts et al., 2015), where the functional interactions of antibodies are harder to test due to species differences (Bournazos, Wang, Dahan, Maamary, & Ravetch, 2017). However, they have been implicated in protection from HIV-infection in humans in several settings. In the only partially effective HIV vaccine trial to date, antibodies that mediate ADCC were associated with protection, whereas neutralizing antibodies were not (Haynes et al., 2012). Moreover, in the setting of mother-to-child-transmission, ADCC antibodies have been associated with both decreased transmission risk and disease progression in infants (Mabuka, Nduati, Odem-Davis, Peterson, & Overbaugh, 2012; Milligan, Richardson, John-Stewart, Nduati, & Overbaugh, 2015). For these reasons, ADCC antibodies, which have the potential to eliminate infected cells, have been increasingly recognized as important to consider for vaccine design and prevention efforts (Lewis et al., 2017). Yet, nothing is known about the process of SHM required for HIV specificity and ADCC function of antibodies.

In light of the relevance of ADCC antibodies in human infections, we deep sequenced the antibody repertoire of an individual who developed potent ADCC antibodies to the HIV envelope (Env) gp120 and gp41 proteins. We used methods that were specifically developed to study antibody evolutionary pathways to infer the naïve precursors and developmental intermediates of six mature ADCC-mediating antibodies that have been previously described (Williams et al., 2015; Williams et al., 2019). Most inferred naïve antibodies did not mediate ADCC. To develop ADCC functionality, inferred naïve antibodies required CDR-localized SHM in one or both chains. In five of six cases, enhancement of ADCC potency to mature levels required mutations in framework regions (FWR), either alone or alongside additional CDR mutations. Interestingly, in one lineage, the developing antibodies first gained the capacity to bind HIV Env and then subsequently acquired ADCC activity due to additional SHM. Overall, binding affinity for Env and facilitation of ADCC were correlated, but we observed cases where binding affinity was similar amongst two antibodies within the same lineage but they differed in their ADCC capacity, suggesting that affinity alone is not sufficient for an antibody to mediate ADCC.

## Results

### Longitudinal sequencing of six ADCC HIV-1 antibody lineages

We previously reported the isolation of six monoclonal antibodies (mAbs), each with potent antibody-dependent cell cytotoxicity (ADCC) activity, from a sample collected 914 days post-infection (D914) from a clade A-infected Kenyan woman (Williams et al., 2015; Williams et al., 2019). All of these mAbs derived from distinct B cell lineages, with four targeting two distinct gp41 Env epitopes (006, 016, 067, 072), and the other two targeting distinct gp120 Env epitopes (105, 157). One of the gp120 targeting mAbs (105) also had modest capacity to neutralize cell-free virus (Williams et al., 2015), whereas the others were non-neutralizing. To infer the ontogeny of these six mAbs, we used next-generation sequencing (NGS) to sequence antibody variable region genes from five longitudinal blood samples collected from the subject, spanning from pre-infection (D-119) to over 4 years post-infection (D1512). Amongst the different antibody chains and timepoints sequenced, the sequencing libraries ranged widely in their sample coverage, between 6-409% with an average of 100% coverage, based on estimated B cell frequencies within the available peripheral blood mononuclear cell (PBMC) samples (Table 1). It is important to note that, despite the name “deep sequencing,” NGS-sampled datasets are relatively shallow compared to the entire repertoire. This is for two reasons: 1) each 10 mL PBMC sample is only ~0.22% of the subject’s whole-body circulating blood volume (~4.5 L), not counting lymphoid organs or peripheral tissue, and 2) circulating human memory B cell repertoires fluctuate over time (Horns, Vollmers, Dekker, & Quake, 2019; Laserson et al., 2014), making it difficult to accurately track clonal B cell families. Despite these limitations inherent to studies of this type, our sequencing efforts successfully returned clonal sequences for all six lineages of interest.

**Table 1.** Longitudinal QA255 antibody variable region sequencing statistics.

| QA255 time point | Live PBMC count (excluding non-viable) | PBMC viability | Ab Chain | Replicate | Raw MiSeq reads | Productive replicate-merged deduplicated sequences | Estimated sequencing coverage within sampled blood PBMCs<sup>b</sup> | Estimated sequencing coverage within QA255 d914 whole-body blood repertoire<sup>c</sup> |
| --- | --- | --- | --- | --- | --- | --- | --- | --- |
| D-119 | 1.39E+07 | 84.22% | IgM | 1 | 486458 | 200254 | 21% | 0.05% |
|  |  |  | IgG | 1 | 604237 | 199570 | 48% | 0.11% |
|  |  |  | IgK | 1 | 705163 | 292543 | 21% | 0.05% |
|  |  |  | IgL | 1 | 406267 | 178221 | 13% | 0.03% |
| D462 | 1.40E+07 | 86.42% | IgG | 1 | 646809 | 504029 | 120% | 0.22% |
|  |  |  |  | 2 | 923103 |  |  |  |
|  |  |  | IgK | 1 | 57096 | 506989 | 36% | 0.08% |
|  |  |  |  | 2 | 967693 |  |  |  |
|  |  |  | IgL | 1 | 201566 | 402456 | 29% | 0.06% |
|  |  |  |  | 2 | 586568 |  |  |  |
| D791 | 3.90E+06 | 97.50% | IgG | 1 | 126955 | 263848 | 226% | 0.22% |
|  |  |  |  | 2 | 776578 |  |  |  |
|  |  |  |  | 3<sup>a</sup> | 522308 |  |  |  |
|  |  |  | IgK | 1 | 527031 | 568736 | 146% | 0.22% |
|  |  |  |  | 2 | 934780 |  |  |  |
|  |  |  |  | 3<sup>a</sup> | 1057984 |  |  |  |
|  |  |  | IgL | 1 | 47368 | 212922 | 55% | 0.12% |
|  |  |  |  | 2 | 387730 |  |  |  |
|  |  |  |  | 3<sup>a</sup> | 537611 |  |  |  |
| D1174 | 1.80E+06 | 100.00% | IgG | 1 | 497094 | 220740 | 409% | 0.22% |
|  |  |  |  | 2 | 665981 |  |  |  |
|  |  |  |  | 3<sup>a</sup> | 678973 |  |  |  |
|  |  |  | IgK | 1 | 254890 | 415601 | 231% | 0.22% |
|  |  |  |  | 2 | 797239 |  |  |  |
|  |  |  |  | 3<sup>a</sup> | 929145 |  |  |  |
|  |  |  | IgL | 1 | 228992 | 307415 | 171% | 0.22% |
|  |  |  |  | 2 | 453935 |  |  |  |
|  |  |  |  | 3<sup>a</sup> | 510711 |  |  |  |
| D1512 | 1.00E+07 | 79.37% | IgG | 1 | 370198 | 155262 | 52% | 0.12% |
|  |  |  | IgK | 1 | 408498 | 214651 | 21% | 0.05% |
|  |  |  | IgL | 1 | 93848 | 55785 | 6% | 0.01% |
|  |  |  |  |  | averages: | 293689 | 100% | 0.13% |

<sup>a</sup> Library replicate performed using unique molecular identifiers during cDNA synthesis step.
<sup>b</sup> Sequencing coverage was calculated for PBMCs using QA255 D914 B cell frequency statistics from the cell sort (Williams et al., 2015).
<sup>c</sup> Whole-body sequencing coverage was calculated assuming 10 ml of blood was sampled from a total of 4500 ml adult blood volume.

We identified sequences clonally-related to the gamma heavy (VH) and lambda (VL) or kappa (VK) light chains of the six mature mAbs (Table 2; Table 2-figure supplement 1A-B), inferred the most likely naive ancestors of each mAb chain, annotated clonal family V, (D), and J gene usage (Table 2), and calculated the degree of mutation in each lineage over time (Table 2; Table 2-figure supplement 1C). Overall, the level of SHM in the mature D914 mAbs ranged from 3.3-11.5%, similar to our previous reporting of these lineages using different annotation methods (Williams et al., 2015; Williams et al., 2019). Incidentally, consensus among clonal sequences within families revealed potentially artifactual cloning mutations at the 5’ and 3’ ends of the variable regions in some of the previously characterized mature mAbs (Williams et al., 2015; Williams et al., 2019) and suggested, overall, that the consensus sequences were the most likely sequences for the antibodies. When directly compared, the sequence-corrected mAbs demonstrated equivalent epitope binding strength and ADCC activity to those previously studied (data not shown). Thus, we corrected the 5’ and 3’ ends of the mature D914 mAb variable regions to consensus sequences among each clonal family for use in this study (Supplementary File 1).

**Table 2.**
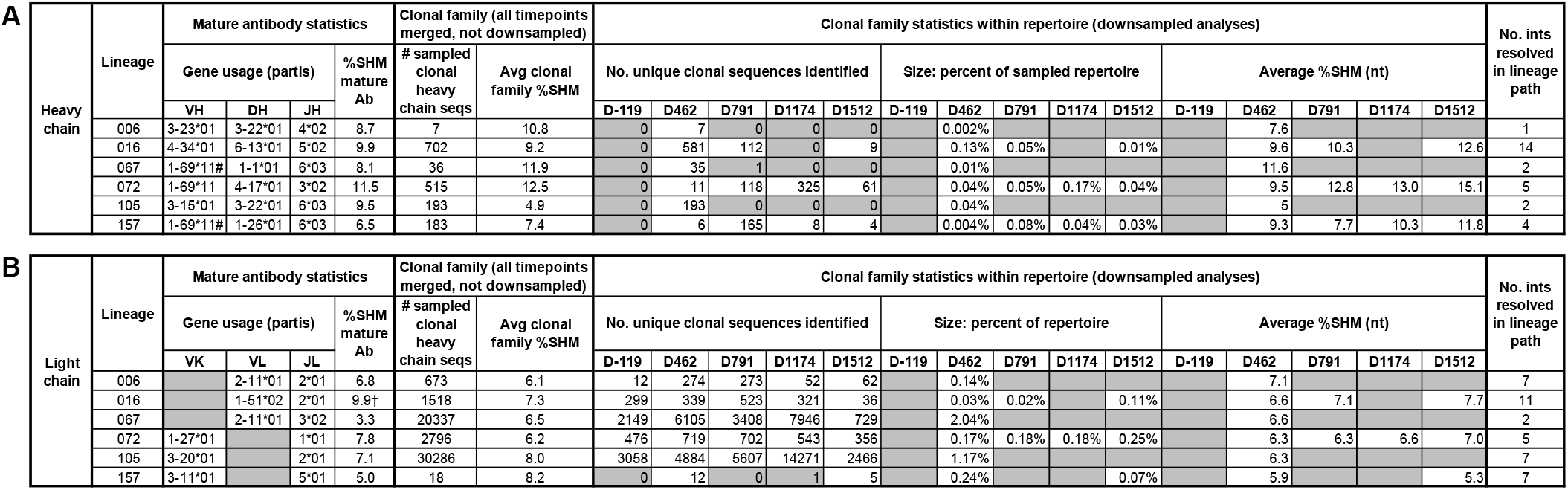
Clonal sequence analyses for ADCC antibody lineages. (A, B) Characteristics and statistics of heavy (A) and light (B) chain clonal families. Percent SHM was calculated as the mutation frequency at the nucleotide level compared to the predicted naïve allele, as determined by the per subject germline inference for QA255. Statistics calculated for individual timepoints within the context of the full subject repertoire were downsampled to 50-150K sequences for computational manageability. Light chain clonal family size statistics are reported but unreliable due to overclustering caveats that are explained in the methods. Per-timepoint clonal family statistics were excluded if ≤1 clonal heavy or light chain sequences were identified. #: The VH gene used in 067 and 157 lineages was determined to be an allele not cataloged in IMGT^®^ (www.imgt.org): 1-69*11+A147G+C169Tl; the 157 lineage VK gene was also a new allele: 3-11 *01+T5A+T9A+T36G+G84A. †: Percent SHM for 016 light chain lineage includes a 3-nt insertion in the CDRL3. SHM: somatic hypermutation; ints: intermediates. Longitudinal values for “No. of unique clonal sequences identified” are plotted for heavy and light chains in Table 2-figure supplement 1, along with longitudinal values for “Average %SHM (nt)” for heavy chains. Graphics displaying the most probable routes of antibody lineage maturation corresponding to the “No. of ints resolved in lineage path” are available in Table 2-figure supplement 2.

### mAbs constructed from D462 NGS sequences mediated ADCC

Each lineage varied in the time point(s) from which we identified clonal sequences, with most clonal VH sequences existing in the D462 datasets for four of the six lineages (Table 2A). The NGS sequences within each lineage that shared the highest nucleotide (nt) identity with the D914 mature VH variable regions, ranging from 88-99%, were found within the D462 or D791 timepoints (Supplementary File 2). As test cases, we synthesized antibodies with VH sequences derived from the D462 NGS results that were similar to the mature 006 and 016 mAbs (88.5% and 97.5% nt identity, respectively). The 006-NGS_VH_ was paired with 006-NGS_VL_ light chain that had 95.8% nt identity to the mature 006 VL (Figure 1-figure supplement 1A) and, since this option was not available for 016 VL, the 016-NGS_VH_ was paired with a computationally-inferred light chain (016-2_VL_) that preceded the mature in development (Figure 1-figure supplement 1B). Both NGS-based mAbs demonstrated full ADCC function (Figure 1), suggesting that ADCC function likely developed within these lineages by D462 post infection.

**Figure 1.**
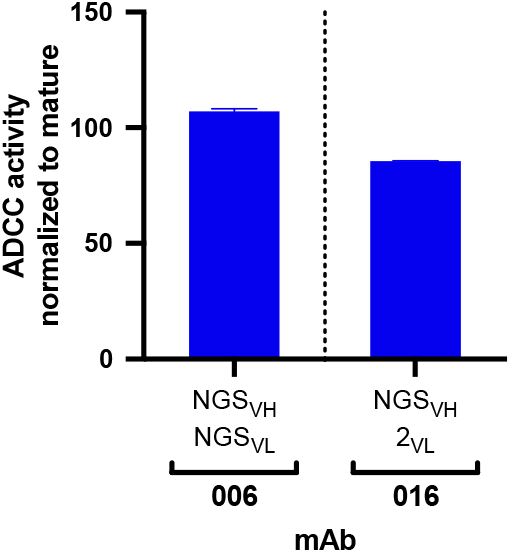
ADCC activity in antibodies made from NGS-sampled sequences. RFADCC activity of antibodies that use NGS-sampled heavy chains that are most similar in sequence to the mature 006 or 016 antibodies. The 006-NGSVH was paired with an NGS-sampled light chain; the 016-NGSVH was paired with a computationally-inferred mature-like sequence 016-2VL. Data reflect two independent experiments.

### Inferred naïve mAbs vary in antigen binding capability and ADCC function

Inferred naïve ancestor mAbs across lineages varied in their ability to bind HIV Env and facilitate ADCC function. Of the gp41-targeted lineages, only the inferred naïve precursor of the 072 lineage bound gp41 (K_D_ = 34.4 nM) and mediated detectable ADCC, although the activity was very low (Figure 2A,C). Both naïve mAbs from the gp120-targeted lineages bound monomeric gp120 (K_D_ = 23.4 nM for 105-0_VH_0_VK_ and 41.1 μM for 157-0_VH_0_VK_; Figure 2B). Interestingly, despite the 105 naïve mAb having higher binding affinity compared to 157 naïve mAb, it did not mediate ADCC, while the 157 naïve mAb mediated potent ADCC (Figure 2C).

**Figure 2.**
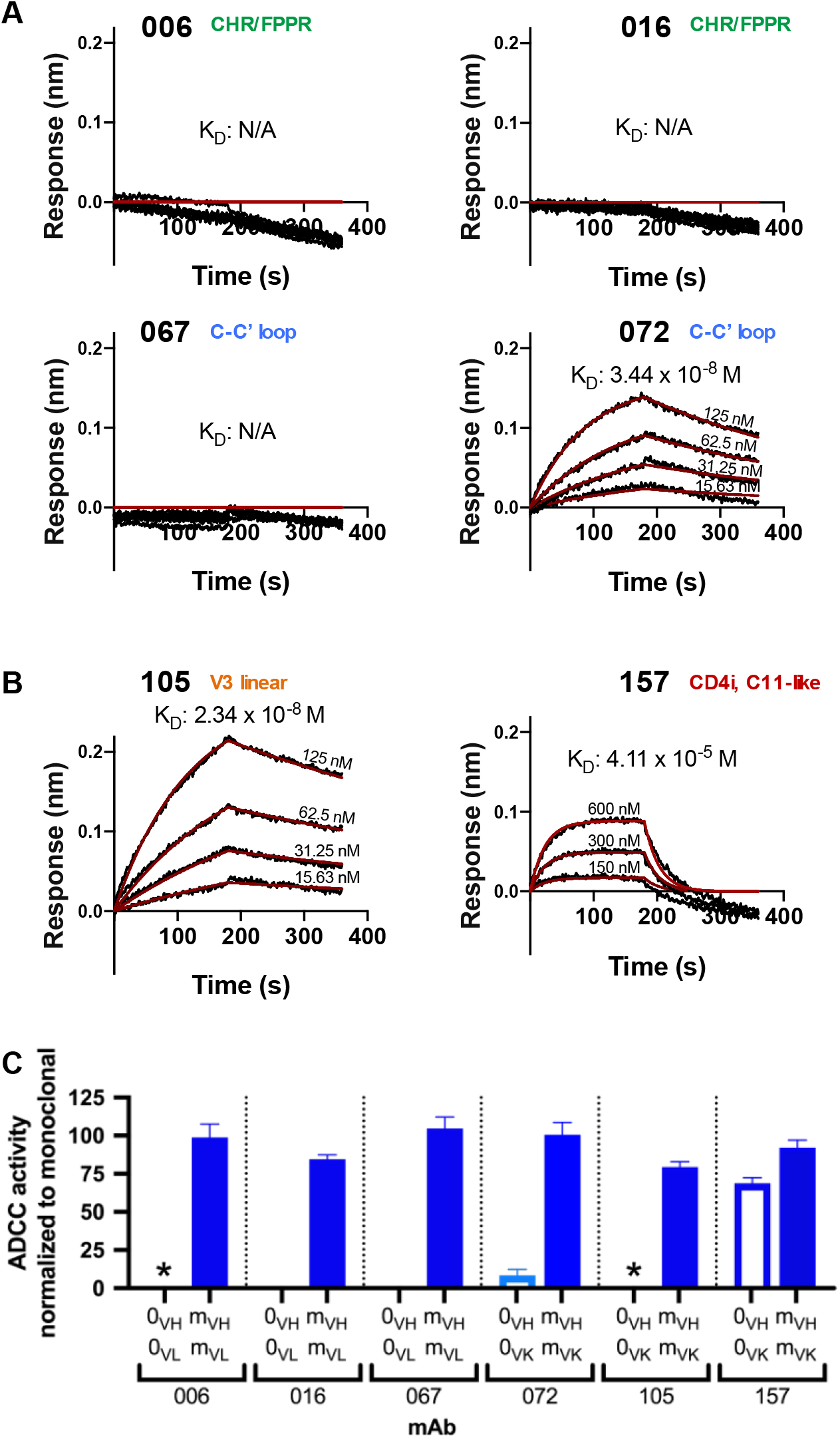
Inferred naïve mAbs from six lineages vary in antigen binding capability and ADCC function. (A-B) Binding kinetics of the inferred naïve antibody (ligand) from each lineage to indicated concentrations of monomeric C.ZA.1197MB gp41 ectodomain (A) or BL035.W6M.C1 gp120 (B) (analyte). Best fitting lines (red) to a 1:1 binding model of ligand:analyte binding are shown. Data are representative of two independent experiments. (C) Positive control-normalized RFADCC activity of inferred naïve antibodies compared to their respective mature antibodies. Normalization is described in Methods. Data are represented as mean ± SEM and reflect at least five independent experiments. 0: naïve; m: mature. See also Figure 2-figure supplement 1.

Typically, the uncertainty in the computational inference of antibody lineage naïve precursors is largely ignored, even though single amino acid differences in the naïve mAb can result in differences in HIV Env binding and functional properties (Yuan, Li, & Zhang, 2011). To account for inference uncertainty, we also tested “alternative naive” mAbs for each lineage that were inferred not by partis (Ralph & Matsen, 2016b), but by linearham (Dhar, Ralph, Minin, & Matsen, 2020), a second computational method that jointly models V(D)J recombination and evolutionary history. For 7 of 12 antibody chains, this alternative approach generated naïve sequences that differed from the original naïve sequences by one to four amino acids within the CDR3 and/or FWR4 regions (Supplementary File 1). For the remaining five, the alternative method inferred the same naïve sequence as the original method. When tested, the alternative naïve sequences did not alter antigen binding or function in five of six antibody lineages when compared to the original naïve mAbs (Figure 2-figure supplement 1A). However, in the 157 lineage we found that the alternative naïve VK chain (157-A_VK_) differed from the original naïve (157-0_VK_) by two adjacent amino acids in the CDRL3 (Figure 2-figure supplement 1D). When 157-A_VK_ was paired with either the original (157-0_VH_) or alternative (157-A_VH_) VH chains, it abolished gp120 binding and ADCC functionality (Figure 2-figure supplement 1B,C). Thus, there is some uncertainty about whether the true 157 lineage naïve precursor was specific for HIV and/or capable of mediating ADCC.

### Characterization of lineage intermediates that confer HIV specificity and ADCC function

To reconstruct probable developmental routes that led to the mature D914 mAb heavy and light chains, we used Bayesian phylogenetic lineage inference, which emphasizes only the inferred ancestral intermediates that have high relative statistical confidence (Simonich et al., 2019). Among the twelve heavy and light chain lineages for the six mAbs, we resolved anywhere from 1 to 14 probable intermediate sequences that lay between the naïve and the mature sequences (Table 2, Table 2-figure supplement 2). Of note, 8 of 12 of thesecomputationally-inferred lineages were validated by the existence of NGS-sampled sequences that were identical either in nucleotide or amino acid sequence to at least one of the inferred intermediates (Table 2-figure supplement 2).

To focus our subsequent lineage studies on determining which mutations enabled HIV Env binding and ADCC gain-of-function, we employed the following strategy to select inferred intermediates of interest (detailed fully in Methods). First, we chose lineage intermediates with high relative confidence that were near the middle of each inferred lineage for preliminary experiments to determine whether antigen binding or ADCC gain-of-function occurred before or after these “middle” intermediate (Table 2-figure supplement 2). If the middle intermediate had function, we focused on pre-middle inferred intermediates, if available within the computationally-inferred lineages. If the middle intermediate lacked function, we focused on post-middle inferred intermediates. mAbs comprised of select intermediate sequences were tested to define the mutations that conferred HIV binding and ADCC activity in each lineage as follows:

*gp41 antibody 006*: mAb 006 (VH3-23, VL2-11) targets a discontinuous epitope that includes the C-terminal heptad repeat (Williams et al., 2019). Mutations in CDRH2 and CDRH3, totaling 1.4% VH SHM, were required for antigen binding and ADCC activity by 006-1_VH_0_VL_ in the 006 lineage (Figure 3A). Thereafter, six light chain mutations among CDRL1, CDRL3, and FWRL2 augmented gp41 binding and ADCC potency in 006-1_VH_1_VL_.

*gp41 antibody 016*: mAb 016 (VH4-34, VL1-51) targets a similar gp41 epitope as 006 (Williams et al., 2019). Detectable gp41 binding and ADCC function was demonstrated by the 016-1_VH_1_VL_ mAb which contains mutations in heavy and light chain CDRs and FWRs (5.6% VH SHM and 3.6% VL SHM) (Figure 3B). A subsequent CDRL3 insertion of residue S94 allowed for augmented ADCC capacity (016-1_VH_2_VL_). Additional FWR and CDR mutations in the VH chain (016-2_VH_2_VL_) further strengthened gp41 binding and ADCC functionality.

*gp41 antibody 067*: mAb 067 (VH1-69, VL2-11) targets the C-C’ loop (Williams et al., 2019). For the 067 lineage, the 067-0_VH_1_VL_ mAb incorporated a single CDRL3 mutation (0.3% VL SHM) and displayed detectable antigen binding; we could not determine whether there was meaningful ADCC function or not as levels were just above background (Figure 3C). To achieve more convincing ADCC function, two CDRH2 mutations were also required (1.0% VH SHM). Further mutations in either the heavy or light chain FWRs modestly increased ADCC potency to >50% that of the 067 mature mAb.

*gp41 antibody 072*: mAb 072 (VH1-69, VK1-27) targets an overlapping but distinct epitope as 067 (Williams et al., 2019). Unlike the other gp41-targeting lineages, the 072 lineage inferred naïve mAb bound gp41 antigen and mediated weak ADCC, as aforementioned (Figures 2A,C and 3D). To gain ADCC potency, only mutations in the VH chain were necessary; mAbs 072-1_VH_0_VK_ (0.8% VH SHM) and 072-2_VH_0_VK_ (0.5% VH SHM) demonstrate that two substitutions in either the CDRH1 or CDRH2 increased activity, but, when combined in 072-3_VH_0_VK_, they were not additive or synergistic (Figure 3D). Instead, FWRH3 and FWRH4 mutations in mAb 072-4_VH_0_VK_ were required for ADCC activity comparable to that of 072 mature mAb.

*gp120 antibody 105*: mAb 105 (VH3-15, VK3-20) targets a V3 linear epitope (Williams et al., 2015). The inferred 105 lineage naïve and 105-0_VH_1_VK_ mAbs bound gp120 but displayed only indeterminate levels of ADCC activity, as defined in Methods (Figure 4A). 105-1_VH_1_VK_ mAb demonstrates that heavy chain CDRH1 and CDRH2 mutations (1.9% VH SHM) were required for ADCC gain of function, even though these mutations did not affect binding affinity (Figure 4A). We note that the FWR4 mutation in the 105-1_VH_ chain was likely a computational artifact and unlikely to affect mAb function, as this mutation is absent in the mature 105 mAb.

*gp120 antibody 157*: mAb 157 (VH1-69, VK3-11) targets a CD4-induced, C11-like epitope (Williams et al., 2015). As detailed in Figure 2-figure supplement 1 and further shown in Figure 4B, inferred naives 157-0_VH_A_VK_ and 157-0_VH_0_VK_ differed in their binding and ADCC capabilities, leaving us uncertain about whether this lineage was able to bind gp120 and facilitate ADCC since inception or gained these capacities following two CDRL3 mutations (Figure 4B).

**Figure 3.**
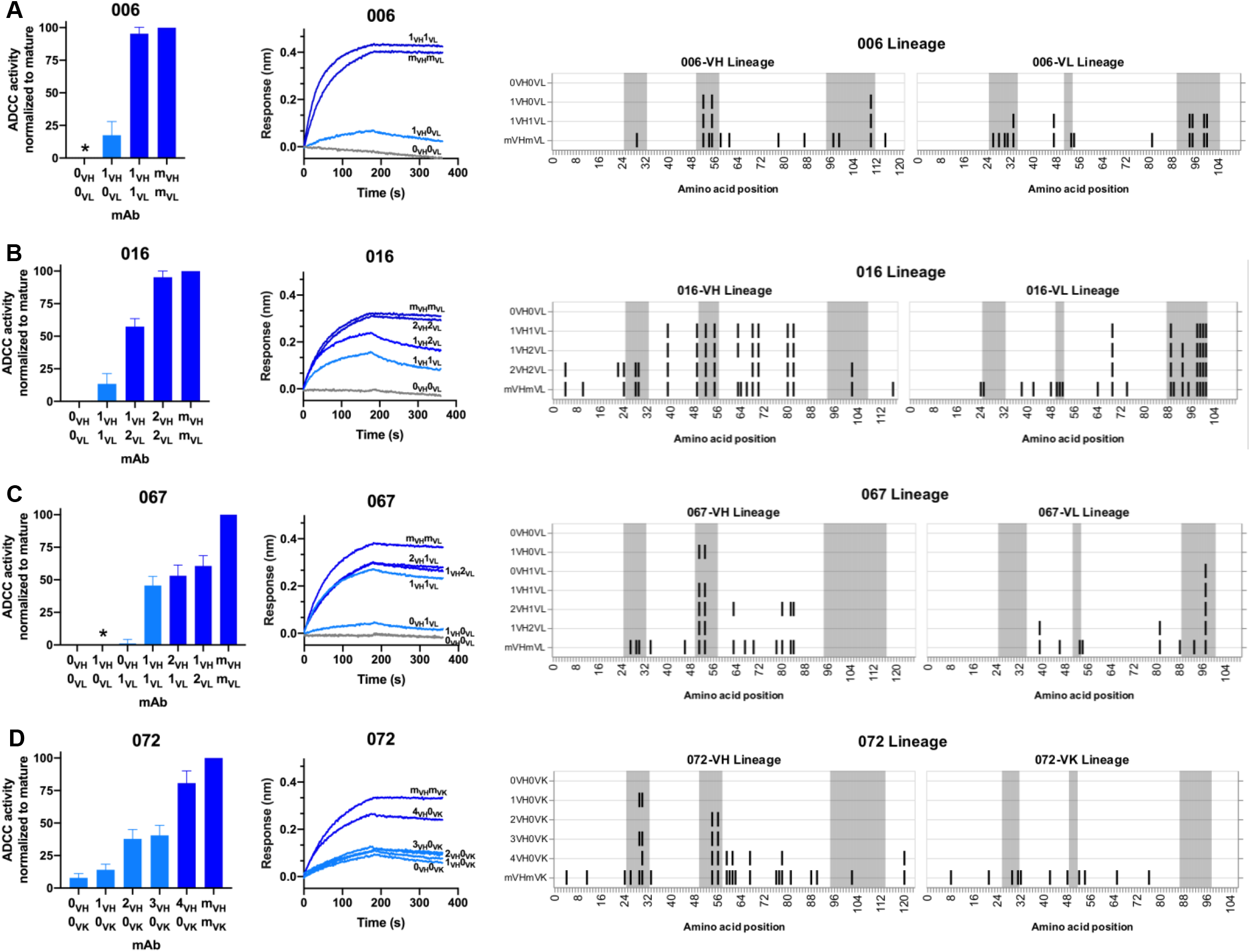
HIV specificity and ADCC development in gp41-targeted antibody lineages. (Left) Mature-normalized RFADCC activity of antibody lineage members for lineages 006 (A), 016 (B), 067 (C), and 072 (D), colored by strength of functionality: no function or indeterminate (gray), low function (<50% of mature activity; light blue), high function (>50% of mature activity; dark blue). Intermediate sequences are numbered consecutively based on their position in the developmental pathway between naïve (0) and mature (m). Asterisks indicate indeterminate activity, as defined in Methods. Data are represented as mean ± SEM and reflect at least four independent experiments, including data presented in Figure 2 for naïve and mature Abs to best account for assay variability and to compare intermediates directly to these antibodies. (Middle) Binding of lineage members (ligand) to monomeric C.ZA.1197MB gp41 ectodomain (analyte, 62.5 nM), measured by BLI. Data are representative of two independent experiments. Data are colored based on each antibody’s RFADCC functionality. (Right) mAb heavy and light chain pairings illustrating variable region amino acid substitutions with respect to the reference sequence at the top of each lineage: black lines indicate amino acid substitutions; gray shaded regions demarcate CDRs.

**Figure 4.**
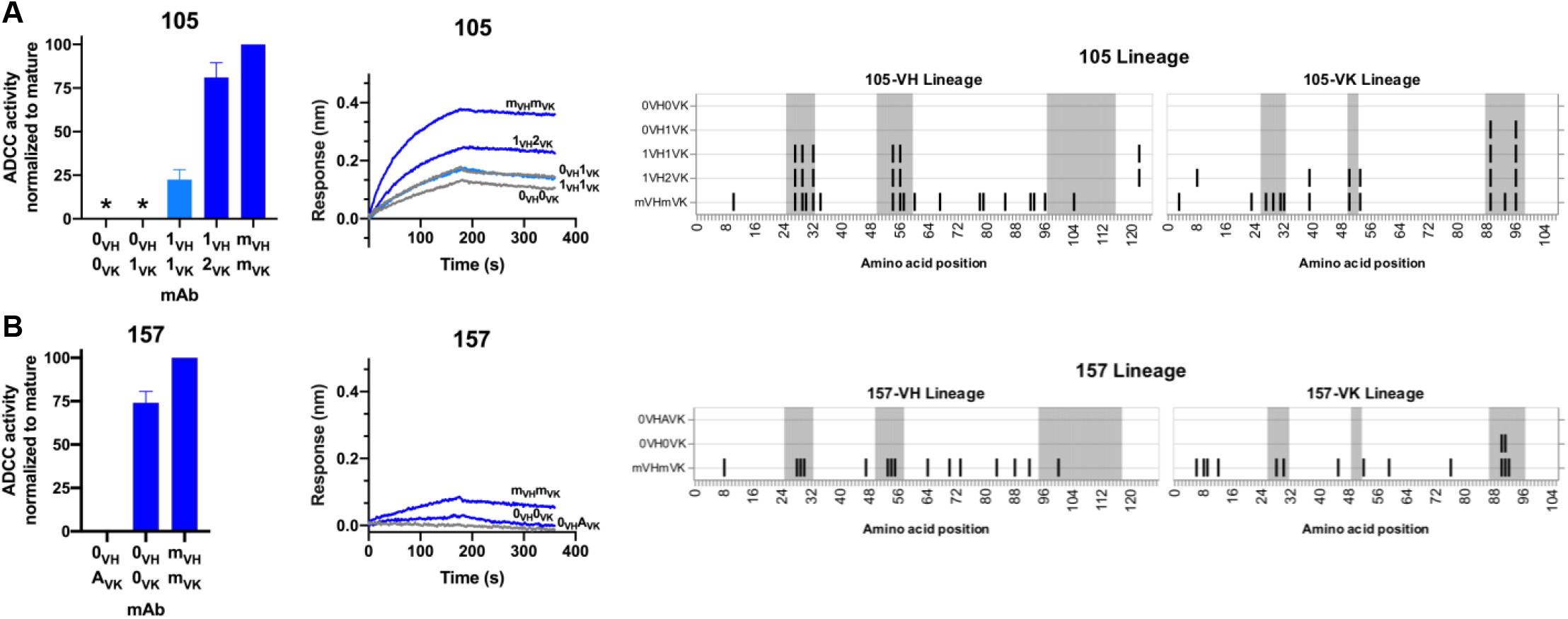
HIV specificity and ADCC development in gp120-targeted antibody lineages. RFADCC activity (left), antigen binding to monomeric BL035.W6M.C1 (analyte, 62.5 nM) as measured by BLI (middle), and mAb heavy and light chain pairings (right) for the 105 and 157 lineages all displayed as in Figure 3. Asterisks indicate indeterminate activity, as defined in Methods. RFADCC data are represented as mean ± SEM and reflect at least four independent experiments, including data presented in Figure 2 for naïve and mature Abs to best account for assay variability.

### Themes in ADCC development pathways among six lineages

Overall, the majority of lineages, regardless of epitope specificity, ultimately required substitutions in both the heavy and light chains to develop ADCC function >50% as potent as their respective mature mAb (Figure 5). In five of six lineages, detectable binding accompanied ADCC capacity, with lineage 105 being the exception that bound prior to developing ADCC function. Most notably, substitutions in CDRs were typically required for ADCC gain-of-function, while subsequent FWR substitutions, either alone or alongside additional CDR mutations, augmented this activity (Figure 5). A graphical summary of the key developmental steps for each lineage is presented in Figure 5-figure supplement 1.

**Figure 5.**
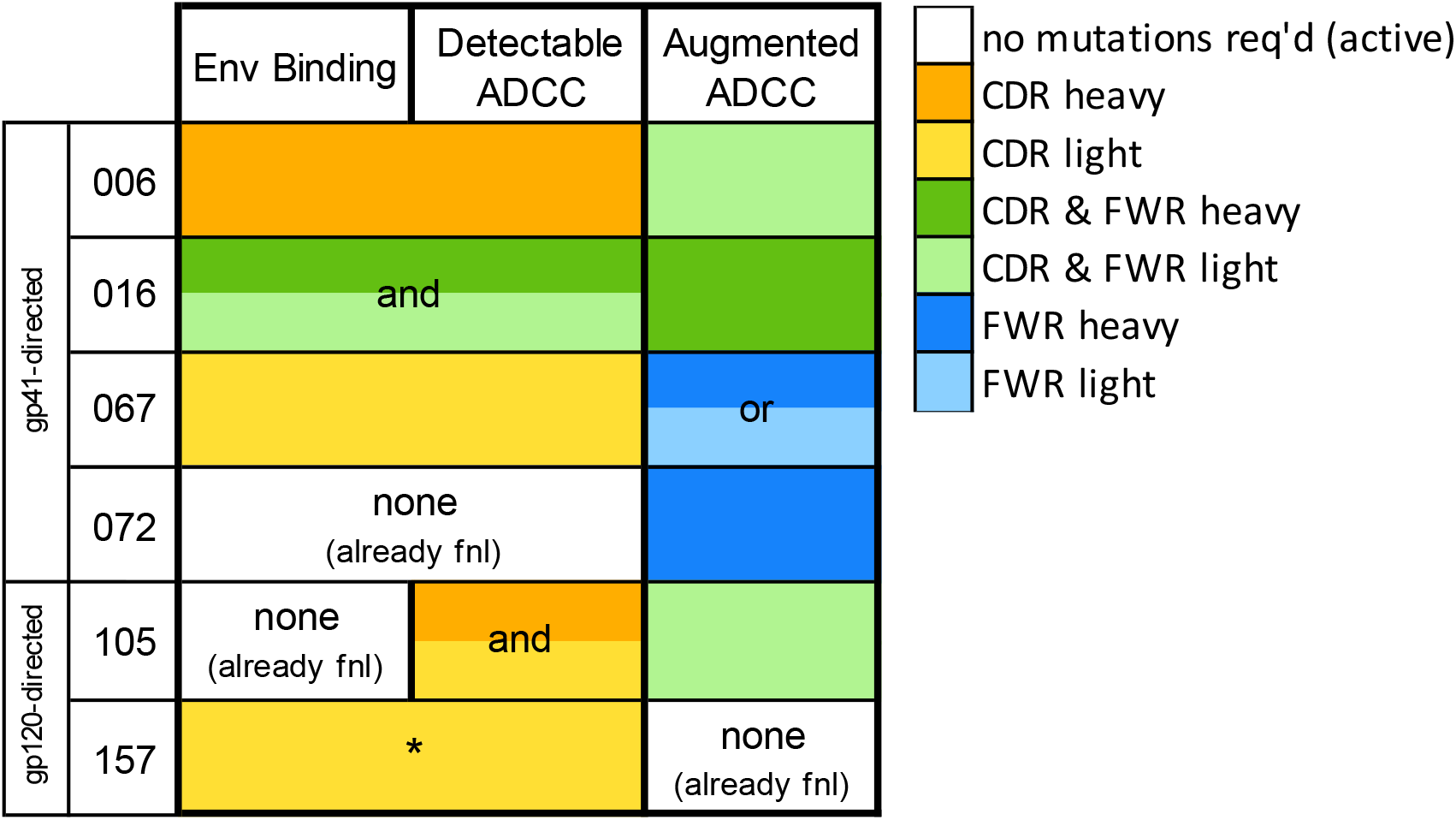
Locality of mutations required for gain of function. Summary of the location of mutations critical for acquiring binding and activity functions in all five ADCC Ab lineages, respective to the previous step, i.e. building upon one another to gain functions. Lineages are summarized by row. Detectable ADCC function excludes lineage members with indeterminate ADCC activity, as defined in Methods. Augmented ADCC activity indicates lineage members with activity most comparable to that of their respective mature. *157-0_VH_A_VK_ required mutations for function while 157-0_VH_0_VK_ did not; Env binding and detectable ADCC determinants were defined based on this comparison. fnl: functional.

### Antigen binding affinity and ADCC function correlate

In our detailed analyses of binding and ADCC activity, there were three lineages (067, 072, and 105) where we observed two mAbs from the same lineage having similar binding affinities but different ADCC capabilities. To visualize the relationship between binding affinity and ADCC potency, we plotted mAb binding affinities (K_D_) against corresponding RFADCC potencies and found that these functionalities were positively correlated in all lineages (Figure 6). It is notable that the antibodies in the 157 lineage that mediate potent ADCC (naïve 157-0_VH_0_VK_ and mature 157-m_VH_M_VK_) bound gp120 antigen with much lower affinity than antibodies with similar ADCC potency in the other five gp41- or gp120-targeted lineages. These data indicate that binding affinity alone does not dictate ADCC potency.

**Figure 6.**
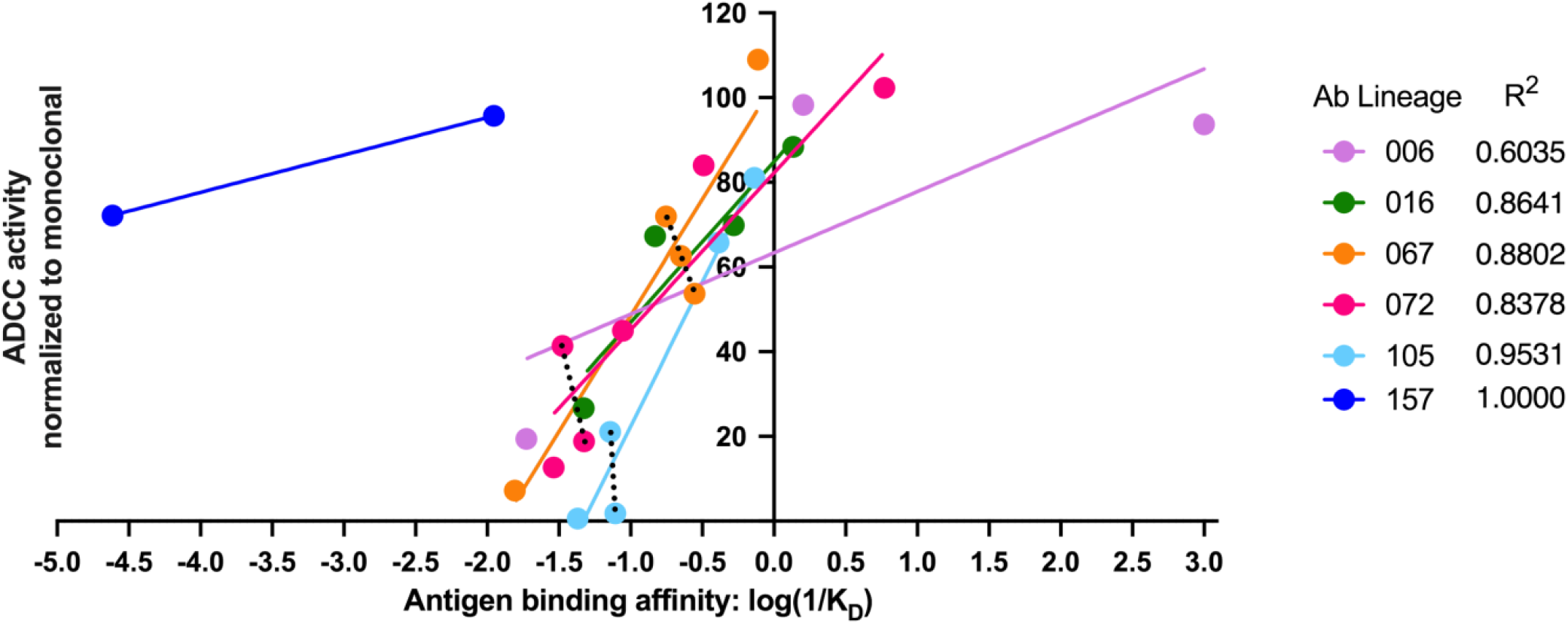
Antigen binding affinity and ADCC function correlate. For antibodies within lineages 006, 016, 067, 072, and 105 (indicated by different colors), the log of reciprocal binding affinity, as determined by representative BLI data of least two independent experiments, is plotted against positive control-normalized RFADCC activity (the data shown in Figures 3 and 4). Antibodies that did not detectably bind antigen by BLI are excluded as none of these mediated ADCC. Dotted lines highlight comparisons between antibodies within the same lineage that have similar binding affinities, but different RFADCC activities.

## Discussion

Renewed interest in ADCC-capable antibodies as important to HIV vaccine responses has highlighted the need for a better understanding of their natural development (Forthal & Finzi, 2018). Here we report the evolution of six ADCC antibody lineages within a single HIV-infected individual. The inferred naïve precursors of these ADCC lineages varied in their abilities to bind HIV antigen, a finding that agrees with studies of HIV-neutralizing antibody lineages (Stamatatos, Pancera, & McGuire, 2017). To achieve potent ADCC activity, most lineages required mutations in both their heavy and light chains. Generally, ADCC function was first achieved through mutations in CDRs, while increased ADCC potency required additional mutations in FWR. ADCC activity accompanied antigen binding in all lineages except one; in the exception the V3/gp120-targeting 105 lineage demonstrated binding capacity prior to developing ADCC functionality. Although there was some evidence that binding affinity was not solely responsible for ADCC capability or potency, binding affinity and ADCC activity were largely correlated. In sum, this study presents six examples of developmental pathways taken by ADCC-mediating antibodies, highlighting that there are common themes in the ontogeny of HIV-specific ADCC antibodies, but that each evolutionary pathway has unique features.

Although not widely applied, antibody lineage studies require robust inference methods that account for statistical uncertainty if we are to be confident in their conclusions. This is highlighted in this study by the different properties displayed by two very similar inferred naïve precursors of the 157 lineage. The development of the 157 lineage was ambiguous: computational uncertainty in two CDRL3 residues resulted in multiple probable naïve antibodies with different properties. In these different scenarios, the 157 naïve Ab was either fully capable of both antigen binding and potent ADCC, or it required two CDRL3 substitutions to achieve both functions. In the other five lineages, our use of multiple computational approaches allowed us to report results with high confidence, since the uncertainties present in computational inference of those lineages did not affect functional characteristics.

Three gp41-specific antibody lineages were not capable of binding HIV Env by their inferred naïve precursors, as measured by BLI; they developed this ability through mutation. The lack of detectable antigen binding by inferred naïve precursors is not altogether surprising as this has been observed for a number of HIV bnAbs (Stamatatos et al., 2017). Binding capacity was gained in two of the gp41-directed lineages (006 and 067) through limited mutations localized in the CDRs of single antibody chains (either heavy or light, respectively). The other gp41-directed lineage, 016, which targets a similar epitope as the 006 lineage, required many mutations amongst the CDRs and FWRs of both chains, highlighting that the gain of activity by different lineages can vary even if they are targeting the same epitope. As is true for all studies of this type, it is possible that our methods were not sensitive enough to detect low level binding or that, despite our best efforts, the inferred naïve sequences were inaccurate. Gp41-directed antibodies have been found to recognize self-antigens (Verkoczy & Diaz, 2014; Verkoczy, Kelsoe, & Haynes, 2014) and this, or its response to another antigen, could explain how the 016 lineage emerged before gaining HIV-specificity.

We chose the RFADCC assay because RFADCC signal has been repeatedly correlated with disease outcome in humans (Dhande et al., 2018; Lewis et al., 2019; Mabuka et al., 2012; Madhavi et al., 2014; Milligan et al., 2015; Ruiz et al., 2017), thus making it relevant to understanding the development of potentially protective responses. There was evidence that the 072 lineage had ADCC function from inception, although the amount of ADCC was low. The remaining lineages required an average of 1.2% SHM to gain detectable ADCC function (ranging 0.1-4.3% SHM). For augmentation of ADCC function to >50% of mature activity, the lineages required an average of 2.2% SHM (range 0.6-4.8% SHM). NGS-sampled sequences that were present at 462 dpi displayed high function with 5.3% SHM (006 lineage). Low mutation levels to achieve potent ADCC are promising for ADCC vaccine efforts, especially since these six characterized lineages have desirable characteristics for combating HIV, such as potent ADCC activity, cross-clade ADCC breadth, highly conserved epitopes, and potential to help clear infected cells (Williams et al., 2015; Williams et al., 2019). Moreover, these levels of mutation are easily attainable by vaccines (Schramm & Douek, 2018).

Importantly, most lineages required FWR mutations in one or both chains to achieve potent ADCC functionality, with or without additional CDR mutations. Though they can cost thermostability (Henderson et al., 2019), FWR mutations are known to contribute to the properties of HIV-specific bnAbs, predominantly by altering Ab conformation in ways that allow conserved epitopes to be bound with high affinity (Henderson et al., 2019; Kepler & Wiehe, 2017; Ovchinnikov, Louveau, Barton, Karplus, & Chakraborty, 2018). FWR residues contribute to the structure of the Ab and, though they do not usually directly contact the antigen, they can affect binding affinity indirectly (Foote & Winter, 1992). Furthermore, since they indeed affect Ab structure, FWR mutations could confer favorable Ab flexibility or an orientation that mediates better Fc receptor dimerization and ADCC potency (Acharya et al., 2014). FWR mutations in Fab “elbow” regions can affect conformational flexibility and paratope plasticity during bnAb development (Henderson et al., 2019). Here, the 006-1_VH_ lineage intermediate caused ADC gain-of-function and it contained, among other mutations, a CDRH3 elbow mutation Y111F. Future structural studies are warranted to explore how FWR mutations affect Ab structure and function in these lineages.

Binding affinity has been implicated in the regulation of ADCC in several studies on tumor-specific Abs (Mazor et al., 2016; Tang et al., 2007). Here, while binding affinity correlated with ADCC function, it was not solely responsible for ADCC potency. In one lineage, binding capacity preceded detectable ADCC function. Notably, we found that half of the lineages contained clonally-related antibodies with similar binding affinity to one another but markedly different ADCC potencies. Conversely, mAbs within the two gp120-targeted lineages exemplify that there can be dramatic differences in binding affinity among mAbs that mediate equivalent, potent ADCC activity. Thus, ADCC function cannot be predicted based on affinity alone. These data support the notion that factors such as epitope specificity, binding mode, and other antibody-antigen interaction dynamics contribute to ADCC potency, as other studies have emphasized (Acharya et al., 2014; Orlandi et al., 2020; Tolbert et al., 2020).

Our study offers a granular understanding of how ADCC functionality developed in six human Ab lineages. The results of this study are relevant to rational vaccine design, especially as technologies for guiding Ab development continue to emerge through reverse vaccinology efforts (Burton, 2017; Rappuoli, Bottomley, D’Oro, Finco, & De Gregorio, 2016; Schramm & Douek, 2018). Additionally, the reliance on FWR mutations to achieve high ADCC functionality could inform mAb optimization strategies for therapeutic uses.

## Materials and Methods

### Human subject

Peripheral blood mononuclear cell (PBMC) samples were obtained between 1997-2002 from a female HIV-1 seroconverted subject, QA255, who was enrolled in a prospective cohort of HIV-1 negative high-risk women in Mombasa, Kenya (Lavreys et al., 2002). QA255 was 40 years old at the time of HIV infection (D0). Study participants were treated according to Kenyan National Guidelines; QA255 did not receive antiretrovirals at any point during the period in which samples were analyzed for this study. Antiretroviral therapy was offered to all participants in the Mombasa Cohort beginning in March 2004, with support from the President’s Emergency Plan for AIDS Relief. The infecting virus was clade A based on envelope sequence (Bosch, Rainwater, Jaoko, & Overbaugh, 2010). Approval to conduct this study was provided by the ethical review committees of the University of Nairobi Institutional Review Board, the Fred Hutchinson Cancer Research Center Institutional Review Board, and the University of Washington Institutional Review Board. Study participants provided written informed consent prior to enrollment.

### Sample handling and RNA isolation

PBMCs stored in liquid nitrogen were thawed at 37°C, diluted 10-fold in pre-warmed RPMI and centrifuged for 10 min at 300 x *g*. Cells were washed once in phosphate-buffered saline, counted with trypan blue, centrifuged again, and total RNA was extracted from PBMCs using the AllPrep DNA/RNA Mini Kit (Qiagen, Germantown, MD), according to the manufacturer’s recommended protocol. RNA was stored at −80°C.

### Sequencing of antibody gene variable regions

Antibody sequencing was performed as previously described (Simonich et al., 2019; Vigdorovich et al., 2016). Library preparation was performed in technical replicate, as indicated in Table 1, using the same RNA isolated from each timepoint: D-119 (119 days prior to HIV infection), D462, D791, D1174, and D1512 post-infection. Briefly, RACE-ready cDNA synthesis was performed using the SMARTer RACE 5’/3’ Kit (Takara Bio USA, Inc., Mountain View, CA) using primers with specificity to IgM, IgG, IgK and IgL, as previously reported (Simonich et al., 2019). One replicate each for D791 and D1174 were prepared using template switch adaptor primers that included unique molecular identifiers in the cDNA synthesis step: SmartNNNa `5’ AAGCAGUGGTAUCAACGCAGAGUNNNNUNNNNUNNNNUCTTrGrGrGrGrG 3’` (Turchaninova et al., 2016), where ‘rG’ indicates (ribonucleoside) guanosine bases. cDNA was diluted in Tricine-EDTA according to the manufacturer’s recommended protocol. First-round Ig-encoding sequence amplification (20 cycles) was performed using Q5 High-Fidelity Master Mix (New England BioLabs, Ipswich, MA) and nested gene-specific primers. Amplicons were directly used as templates for MiSeq adaption by second-round PCR amplification (10-20 cycles), purified and analyzed by gel electrophoresis, and indexed using Nextera XT P5 and P7 index sequences (Illumina, San Diego, CA) for Illumina sequencing according to the manufacturer’s instructions (10 cycles). Gel-purified, indexed libraries were quantitated using the KAPA library quantification kit (Kapa Biosystems, Wilmington, MA) performed on an Applied Biosystems 7500 Fast real-time PCR machine. Libraries were denatured and loaded onto Illumina MiSeq 600-cycle V3 cartridges, according to the manufacturer’s suggested workflow.

### Sequence analysis and naïve inference

Sequences were preprocessed using FLASH, cutadapt, and FASTX-toolkit as previously described (Simonich et al., 2019; Vigdorovich et al., 2016). The sequences from our NGS replicates were merged either cumulatively or on a per-timepoint basis, as appropriate, to achieve the highest depth possible for each analysis. Sequences were then deduplicated and annotated with partis (https://github.com/psathyrella/partis) using default options including per-sample germline inference (Ralph & Matsen, 2016a, 2016b, 2019). Sequences with internal stop codons or CDR3 regions that were out-of-frame or had mutated codons at the start or end were removed. Indel events were identified, tracked, and reversed for alignment purposes. We did not exclude singletons in an attempt to retain very rare or undersampled sequences. Sequencing run statistics are detailed in Table 1. Cumulative and per-timepoint datasets underwent clonal family clustering using both the partis unseeded and seeded clustering methods (Ralph & Matsen, 2016b).

Inference of unmutated common ancestor sequences and simultaneous clonal family clustering was performed using the seed clustering method along with previously-identified QA255 mature ADCC antibody sequences (Williams et al., 2015; Williams et al., 2019) as “seeds”. As an additional measure to ensure highest accuracy for inferred naïve sequences, the uncertainty on each inferred naive sequence was visualized both with the partis --view-alternative-annotations option and by comparing these results to the most likely naive sequences inferred by linearham software (see next section). The comparison revealed only the minor differences that would be expected based on linearham’s method, which uses a more detailed model to infer more accurate naive sequences for individual clonal families. For the unseeded clustering, each dataset was subsampled to 50-150K sequences for computational efficiency. For each dataset, three random subsamples were analyzed and compared to ensure that our subsampling was sufficient to minimize statistical uncertainties. Seeded analyses were not subsampled.

Due to the lack of D genes and much shorter non-templated regions in light-chain rearrangements, computational clustering analyses artifactually overestimate clonality in light chain families, i.e. they cluster together sequences that did not originate from the same rearrangement event, but which come from almost identical naïve rearrangements. This caveat, affecting all unpaired antibody chain sequencing studies, ultimately reduces accuracy in inferring true light chain clonal families and their maturation pathways.

### Alternative naive inference

Alternative naïve sequences were inferred by applying a Bayesian phylogenetic Hidden Markov Model approach using the linearham software (https://github.com/matsengrp/linearham), using the partis-inferred clonal family clusters for each lineage as input (Dhar et al., 2020). Linearham samples naive sequences from their posterior distribution rather than providing a single naive sequence estimate, like other software programs do. The most probable linearham-predicted naïve sequences were compared to the most probable partis-predicted naïve sequences. In families with largely symmetric phylogenetic trees, the two methods return similar results; whereas with highly imbalanced trees, linearham is much more accurate.

### Antibody lineage reconstruction

Antibody lineages were inferred as previously described (Simonich et al., 2019). In order to subsample large clonal families according to phylogenetic relatedness to the antibody chain of interest, initial phylogenetic trees of each QA255 clonal family was inferred using FastTree 2 (Price, Dehal, & Arkin, 2010). This allowed clonal families to be reduced to the 75-100 sequences most relevant to inferring the lineage history of the antibody chain of interest (https://github.com/matsengrp/cft/blob/master/bin/prune.py), which was then analyzed with RevBayes (Höhna et al., 2016) using an unrooted tree model with the general time-reversible (GTR) substitution model. All settings for RevBayes runs were customized to ensure likelihood and estimated sample sizes were >100 for each lineage. MCMC iterations ranged from 10,000-200,000, with thinning frequencies between 10-200 iterations and number of burn-in samples between 10-190. All RevBayes runs were done in technical duplicate (i.e. specifying different starting trees) and duplicates were all confirmed to agree on lineage chronology. RevBayes output was summarized for internal node sequences (https://github.com/matsengrp/ecgtheow, Table 2-figure supplement 2), resulting in summary graphics where relative confidence in unique inferred sequences and amino acid substitutions are represented by color intensity. For each lineage, inferred intermediate sequences found on the most probable lineage paths were selected for study (Supplementary File 1, Table 2-figure supplement 2).

In the 016-VL lineage, we determined that a 3-nt insertion event occurred prior to the inferred_7_712 intermediate within this antibody’s most likely developmental pathway (Table 2-figure supplement 2). Since most insertion-containing clonal NGS sequences (136 of 142) encoded serine at the insertion site (amino acid position 94), we inserted S94 into 016-2VL instead of N94 that would correspond to the mature 016-VL sequence. For thoroughness, the inferred_5_789 016-VL intermediate (which precedes inferred_7_712) was synthesized both without (016-1_VL_) and with (016-2_VL_) the S94 insertion.

### Identification of lineage-like NGS sequences

To determine if the computationally-inferred naïve and ancestor sequences were observed in the NGS data, we performed the following procedure for each lineage (implemented in a script found here: https://git.io/Je7Zp). A local BLAST database was created for each seeded clonal family and queried for sampled sequences that had high nucleotide sequence identity to lineage members using the ‘blastn’ command (Biopython package). E-value of 0.001 was used; other settings were default. Blastn matches for each lineage member were sorted according to their percent nt identity and alignment length. Sampled sequences with 100% nt or 100% aa identity in common with lineage members were noted (Supplementary File 1, Table 2-figure supplement 2). To identify the closest sampled sequences to each VH mature sequence, blastn results per VH mature query were viewed and the highest percent aa identity match was selected.

### ADCC-focused lineage determination

As outlined in our Results, we first selected “middle” lineage intermediates for studying ADCC gain-of-function based on their moderate inclusion of amino acid substitutions (between naïve and mature sequences; Supplementary File 1) and high statistical confidence (Table 2-figure supplement 2), along with, in some cases, their validation by sampled NGS sequences (Table 2-figure supplement 2), and/or concentration of substitutions in complementarity-determining regions (CDRs) (Supplementary File 1). Middle intermediate chains were paired with partner naïve, middle, and mature chains and tested for antigen binding and ADCC function. Based on preliminary results using middle intermediates, ADCC-focused lineages were chosen from remaining pre-middle inferred intermediates or post-middle intermediates depending on whether the middle intermediate showed activity. If multiple relevant intermediate choices were available within the pertinent (early or late) portion of an inferred lineage (Table 2-figure supplement 2), we selected sequences that had amino acid substitutions that were concentrated in CDRs (Supplementary File 1) and, whenever possible, we selected intermediates that were validated by NGS-sampled sequences, as aforementioned (Table 2-figure supplement 2). For 067-VH, 067-VK, 072-VH, and 105-VH lineages, early mutations were implicated in ADCC gain-of-function, but we lacked early lineage resolution and therefore could not select early inferred intermediates to study. Instead, we selectively incorporated CDR substitutions from the middle intermediate sequence into the inferred naïve sequence. Non-conservative FWR3 substitutions were also incorporated into the 067-VH early lineage. For each lineage, intermediate sequences were numbered consecutively based on their position in the developmental pathway between naïve (0) and mature (m). Heavy and light chain pairings for lineage intermediates were based on chronology, which reflected levels of mutation. Each chain was paired with several partner chains for functional assessment.

Ultimately, developmental lineages featuring the minimal number of mAbs to study ADCC development were defined based on levels of mAb mutation and ADCC activity. For each lineage, the minimal antibodies included 1) the inferred naïve, 2) the most mutated intermediate(s) that lacked ADCC activity, if available, 3) the least mutated intermediate(s) that gained detectable ADCC function, and 4) the least mutated intermediate(s) that demonstrated ADCC activity comparable to the mature antibodies (>50%). Mature antibodies were always included as benchmark positive controls.

### Cell lines

#### For antibody production

HEK 293-F cells (RRID:CVCL_D603; originally derived from female human embryonic kidney cells) were obtained from Invitrogen (Thermo Fisher Scientific, Waltham, MA, catalog #R790-07) and grown at 37°C in Freestyle™ 293 Expression Medium (Thermo Fisher Scientific, catalog #12338002) in baffle-bottomed flasks orbiting at 135 rpm. These cells were not further authenticated in our hands.

#### For RFADCC

CEM.NKR cells (RRID:CVCL_X622; originally derived from female human T-lymphoblastoid cells) were obtained from NIH AIDS Reagent Program (Germantown, MD, catalog #458) and grown at 37°C in RPMI 1640 media with added penicillin (100 U/mL), streptomycin (100 μg/mL), Amphotericin B (250 ng/mL), L-glutamine (2mM), and fetal bovine serum (10%) (all from Thermo Fisher Scientific). These cells were not further authenticated in our hands.

### Monoclonal antibody production

Antibody heavy and light chain variable regions were synthesized as FragmentGENES (GENEWIZ^®^, South Plainfield, NJ) and subsequently cloned into corresponding Igγ1, Igκ or Igλ expression vectors (Tiller et al., 2008). Equal ratios of heavy and light chain plasmids were co-transfected into HEK 293F cells using FreeStyle MAX (Thermo Fisher Scientific, catalog #16447100) according to the manufacturer’s instructions. Column-based Pierce™ protein G (Thermo Fisher Scientific, catalog #20397) purification of IgG was done according to the manufacturer’s instructions.

### Rapid and fluorometric ADCC (RFADCC) assay

The RFADCC assay was performed as described (Gomez-Roman et al., 2006; Williams et al., 2019). In short, CEM-NKr cells (NIH AIDS Reagent Program, catalog #458) were double-labeled with PKH-26-cell membrane dye (Sigma-Aldrich, St. Louis, MO) and a cytoplasmic-staining dye (Vybrant CFDA SE Cell Tracer Kit, Thermo Fisher Scientific). The double-labeled cells were coated with clade A gp120 (BL035.W6M.Env.C1, (Wu et al., 2006)) or clade C gp41 ectodomain (C.ZA.1197MB, Immune Technology Corp, New York, NY) (Rousseau et al., 2006) for 1 hr at room temperature at a ratio of 1.5 μg protein: 1 x 10^5^ double-stained target cells. Coated targets were washed once with complete RMPI media (RPMI supplemented with 10% FBS, 4.0 mM Glutamax, and 1% antibiotic-antimycotic, all from Thermo Fisher Scientific). Monoclonal antibodies were diluted in complete RPMI media to a concentration of 100-500 ng/mL and mixed with 5 x 10^3^ coated target cells for 10 min at room temperature. PBMCs (peripheral blood mononuclear cells; Bloodworks Northwest, Seattle, WA) from an HIV-negative donor were added at a ratio of 50 effector cells: 1 target cell. The coated target cells, antibodies, and effector cells were co-cultured for 4 hr at 37°C then fixed in 1% paraformaldehyde (Affymetrix, Santa Clara, CA). Cells were analyzed by flow cytometry (Symphony I/II, BD Biosciences, San Jose, CA) and ADCC activity was defined as the percent of PKH-26+ CFDA-cells after background subtraction, where background (antibody-mediated killing of uncoated cells) was standardized to be 3–5% as analyzed using FlowJo software (FlowJo LLC, Ashland, OR). To mitigate differences in activity observed with different PBMC donors and between experiments, ADCC activity for each sample was normalized to monoclonal positive control mAbs: 167-D (NIH AIDS Reagent Program, Cat #11681) for gp41-targeted Abs and C11 (NIH AIDS Reagent Program, Cat #7374) for gp120-targeted Abs. The activity of an unrelated antibody, FI6v3, that recognizes influenza hemagglutinin protein was used to define the limit of detection. For each replicate experiment, samples were categorized in the following manner: positive (sample > 2*FI6v3), indeterminate (l*FI6v3 ≤ sample ≤ 2*FI6v3), negative (sample < 1*FI6v3). Experiments were excluded if FI6v3 signal was >10% of monoclonal positive control signal. Background (uncoated cells) and negative-control (1*FI6v3) signal was subtracted from each sample’s activity and the resultant values were averaged across experimental replicates, normalized to the respective lineage’s mature mAb activity, and plotted in Prism v8.0c (GraphPad, San Diego, CA). A designation of indeterminate was also assigned in cases where samples were negative in the majority of experimental replicates, but indeterminate or positive in any replicate(s). In such cases, average activity was reported as usual.

### Biolayer Interferometry

QA255 monoclonal antibody binding to monomeric gp120 or gp41 was measured using biolayer interferometry on an Octet RED instrument (ForteBio, Fremont, CA). Antibodies diluted to 8 μg mL^-1^ in a filtered buffer solution of 1X PBS containing 0.1% BSA, 0.005% Tween-20, and 0.02% sodium azide were immobilized onto anti-human IgG Fc capture biosensors (ForteBio). C.ZA.1197MB or BL035.W6M.C1 gp41 was diluted to 250 nM, or as indicated, in the same buffer (above) and a series of up to six, two-fold dilutions were tested as analytes in solution at 30°C. The kinetics of mAb binding were measured as follows: association was monitored for 180 seconds, dissociation was monitored for 180 seconds, and regeneration was performed in 10mM Glycine HCl (pH 1.5). For experiments in which dilution series were run, binding-affinity constants (K_D_; on-rate, K_on_; off-rate, K_dis_) were calculated using ForteBio’s Data Analysis Software 7.0. Responses (nanometer shift) were calculated and background-subtracted using double referencing against the buffer reference signal and non-specific binding of biosensor to analyte. Data were processed by Savitzky-Golay filtering prior to fitting using a 1:1 model of binding kinetics.

### Data and Code Availability

The QA255 longitudinal antibody deep sequencing datasets (Table 1) generated during this study are publicly available at BioProject SRA, accession PRJNA639297 [https://www.ncbi.nlm.nih.gov/bioproject/PRJNA639297/]. The inferred antibody variable region sequences generated in this study have not been deposited in GenBank because computationally-inferred sequences are not accepted, but they are available in Supplementary File 1. Mature antibody GenBank accession numbers are MT791224-MT791235. GenBank accession numbers for HIV Env variant are ABA61538.1 (BL035.W6M.Env.C1 gp120) and AAR27754.1 (C.ZA.1197MB gp41). The custom code generated or used in this study is publicly available on GitHub: prune.py (https://github.com/matsengrp/cft/blob/master/bin/prune.py), ecgtheow (https://github.com/matsengrp/ecgtheow), CFT (https://github.com/matsengrp/cft), and Blast validation (https://git.io/Je7Zp).

### Quantification and Statistical Analysis

Where applicable, raw data were normalized and/or averaged across replicates using Microsoft Excel. Plots were generated using GraphPad Prism version 8.0c. Relevant experimental details, such as use of biological and technical replicates, can be found in figure legends. For RFADCC plots, bars with error bars represent mean and SD. RFADCC experimental exclusion criteria are detailed within the RFADCC method section above.

---

**Table 2-figure supplement 1.**
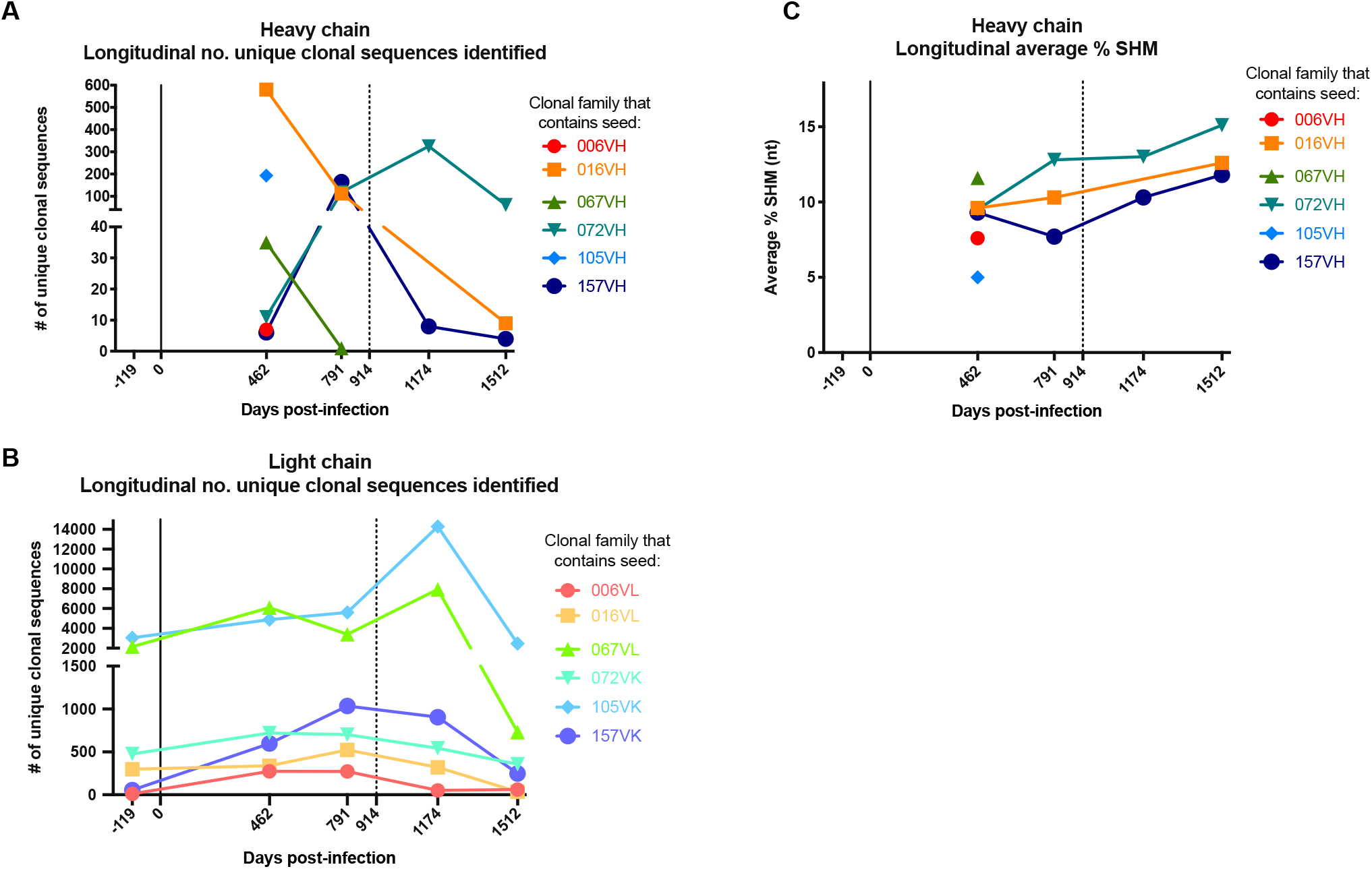
Longitudinal characteristics of heavy and light chain clonal families. (A, B) Number of clonal heavy (A) and light (B) chain sequences identified per sequenced timepoint per lineage. Caveats relating to light chain clusters are described in Methods. (C) Average variable region mutation for each clonal heavy chain family per timepoint. Dashed line at D914 indicates the timepoint of isolation of mature antibodies (Williams et al., 2015; Williams et al., 2019). SHM: somatic hypermutation.

**Table 2-figure supplement 2:**
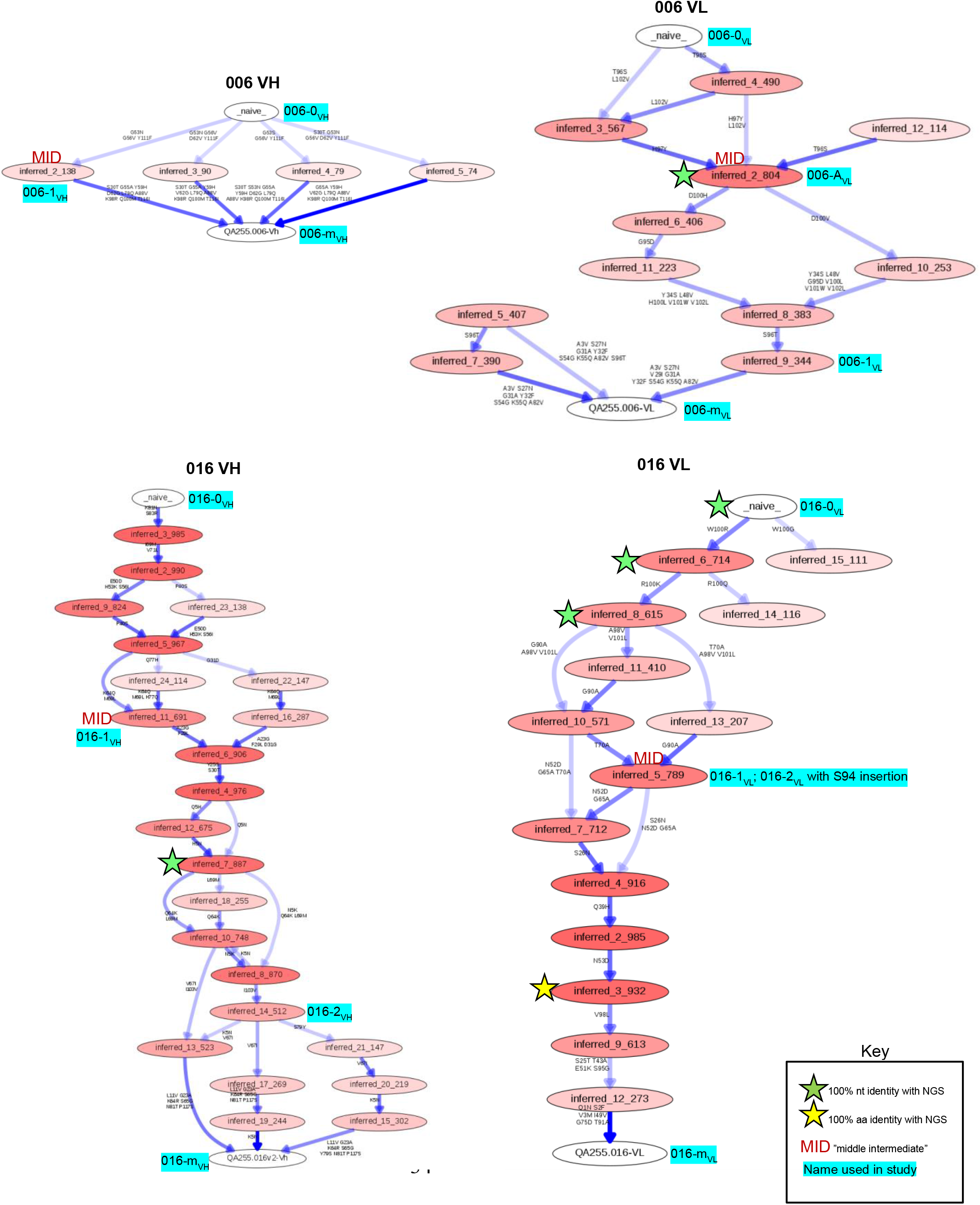
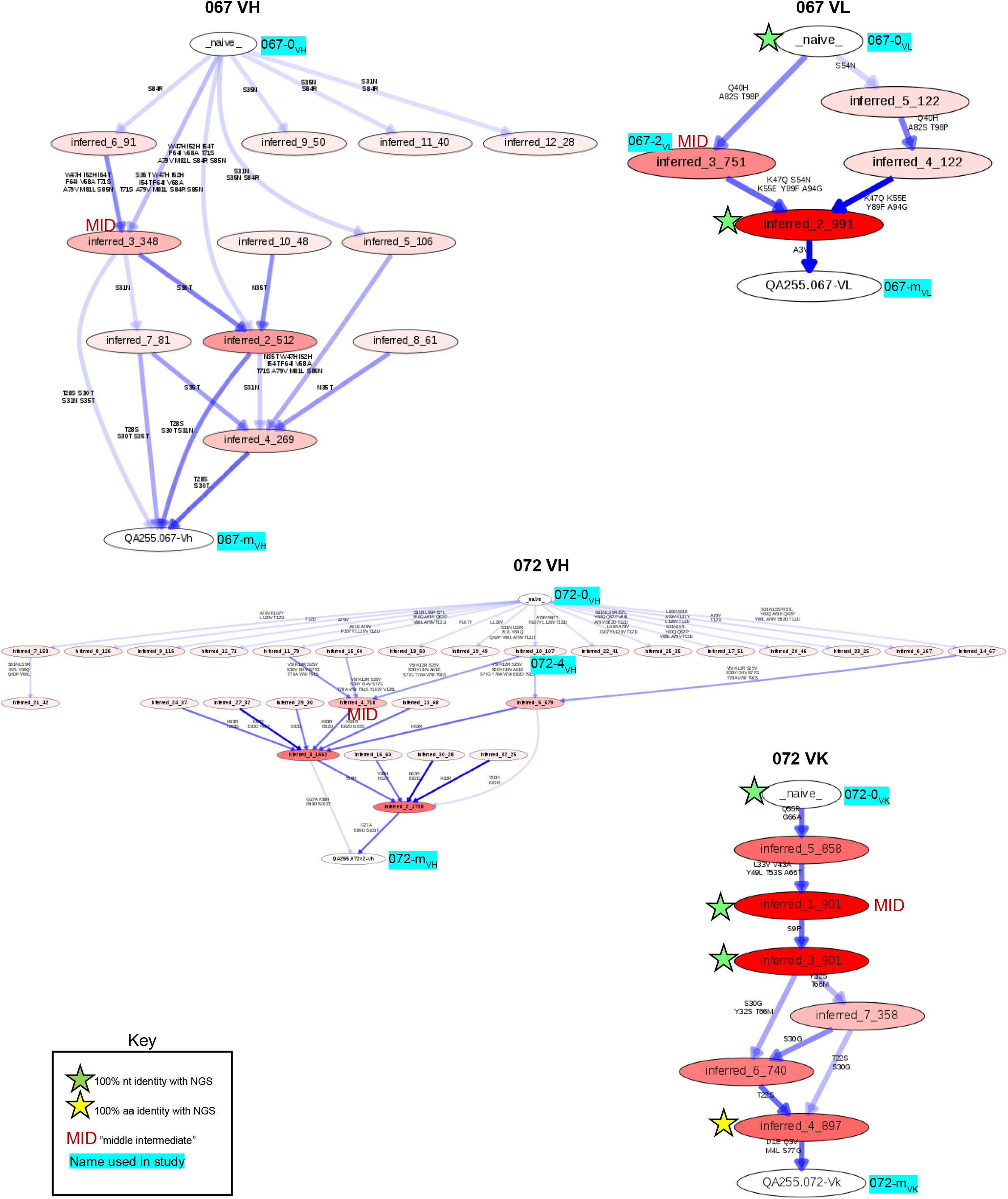
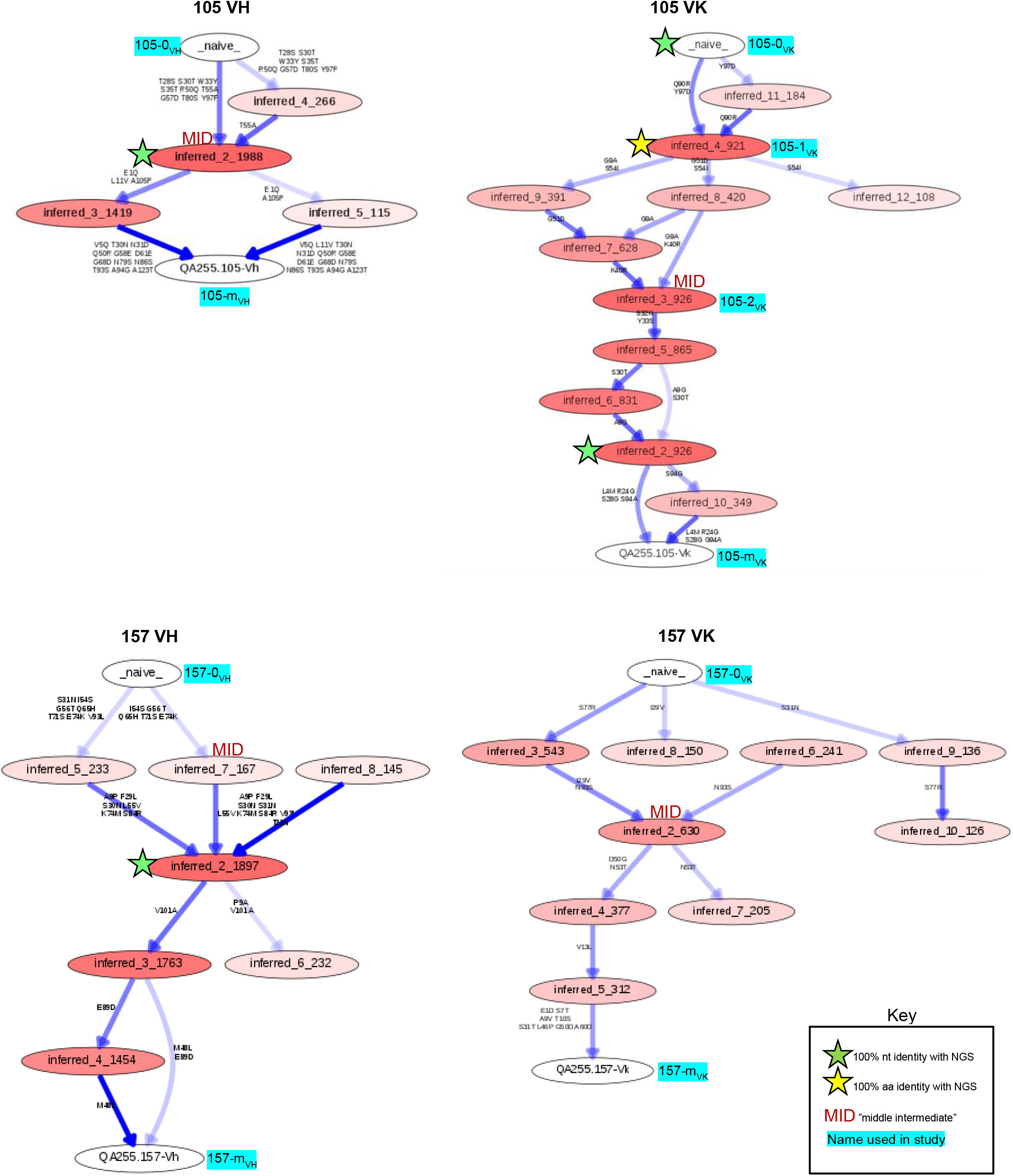
Most probable routes of antibody lineage maturation. Bayesian clonal family phylogenies for each antibody chain were sampled from associated posterior distributions and summarized to display relative confidences for internal node sequences. Resulting graphics display multiple possible lineages of amino acid transitions and their relative confidences for heavy- and lightchain development. Amino acid substitutions (arrows) connect the inferred naive sequence (white node, top) to the mature sequence (white node, bottom) via reconstructed ancestral intermediate sequences (red-shaded nodes). The red shading of nodes is proportional to the posterior probability that this ancestral sequence was present in the lineage. For a given node, the blue shading across arrows arising from that node is proportional to the corresponding transition probability. Low probability nodes were filtered out, resulting in some incomplete pathways within the graphics. Transient mutations are labeled in gray instead of black. Sequences used in functional studies are labeled with their manuscript aliases. Stars denote inferred sequences that were identical in nucleotide (green) or amino acid (yellow) sequence to sampled NGS sequences. “MID” (red) denotes middle intermediate sequence. Labels with cyan highlighting denote inferred sequences that were used in lineage studies. These are incomplete because some lineage members were manually designed (see Methods). If inferred “alternative” naïve sequences matched a lineage member, we included these labels as well for convenience.

**Supplementary File 1. Sequences of six ADCC antibody lineages.** (fasta file) Included are inferred naïve, computationally-inferred lineage intermediate, manually-inferred lineage intermediate, corrected mature, and NGS sequences with high sequence identity to lineage members and/or mature sequences. Sequences are provided as Supplementary File 1 because computationally-inferred sequences cannot be deposited into GenBank.

**Supplementary File 2. Select NGS sequences with highest sequence identity to mature D914 mAb sequences.** (fasta file)

**Figure 1-figure supplement 1.**
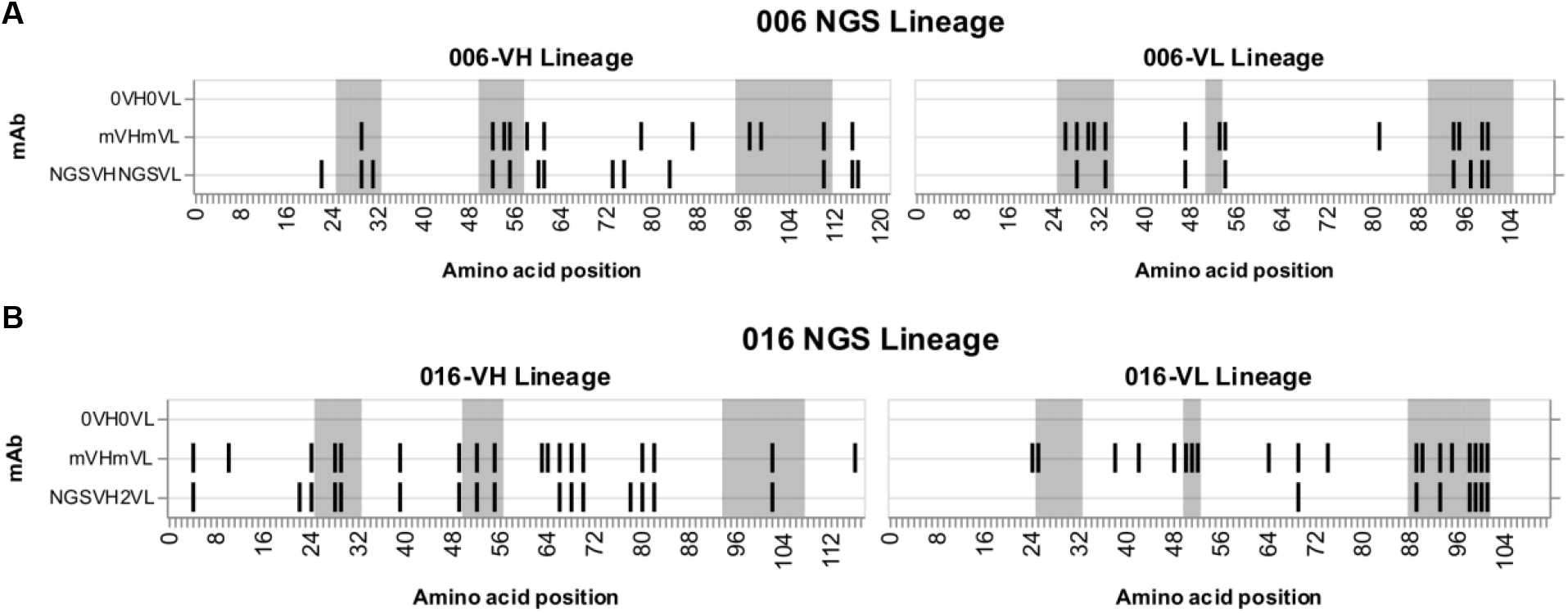
Graphic summaries of antibodies comprised of paired VH and VL NGS-sampled sequences. Graphic summaries of paired VH and VL antibody sequences for 006 (A) and 016 (B) lineages relative to inferred naïve (0) sequences. Shaded regions demarcate CDRs (gray). Vertical black lines indicate variable region amino acid substitutions. m: mature.

**Figure 2-figure supplement 1.**
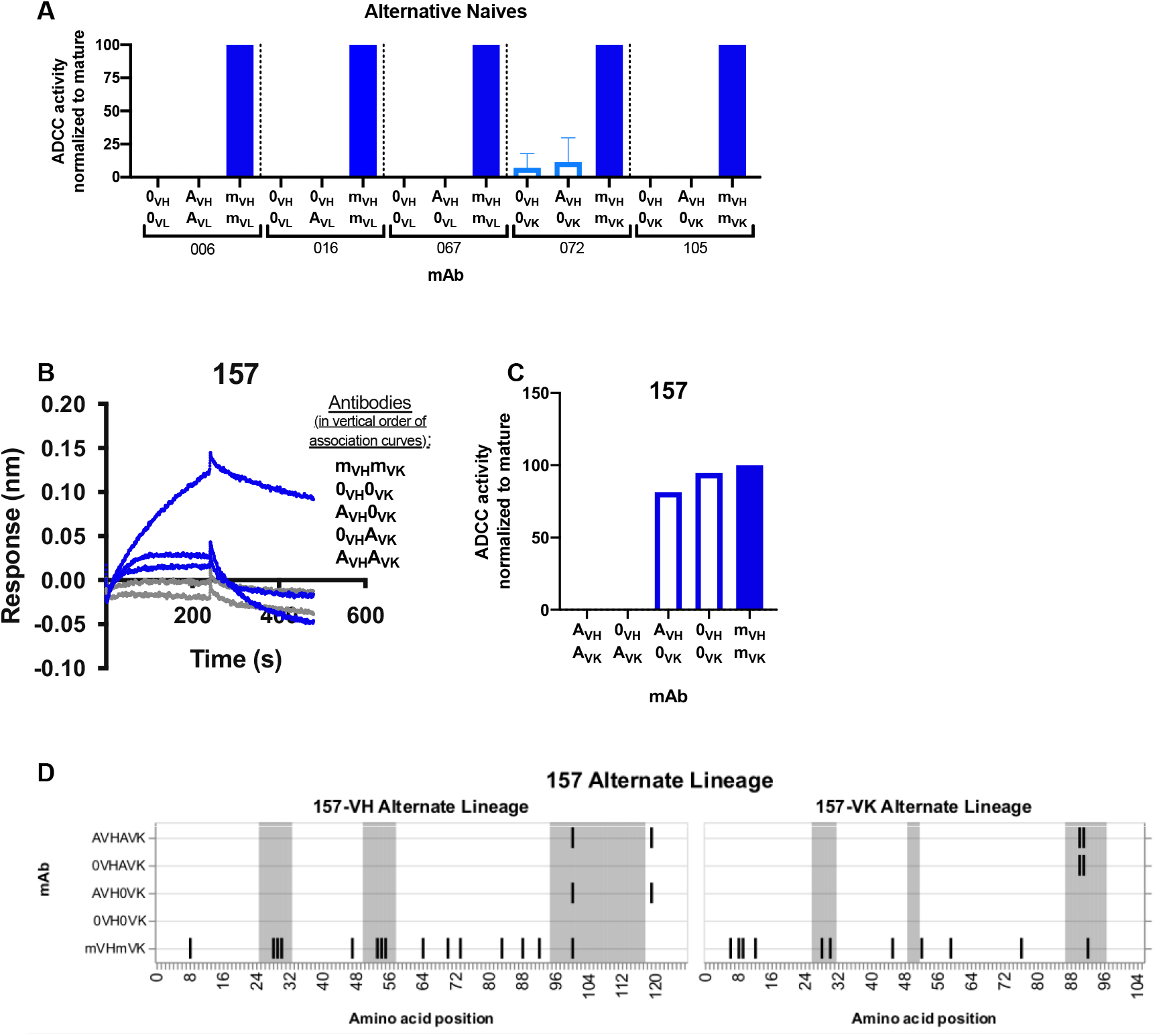
Alternative naïve mAb functionality. (A, C) Functional RFADCC comparisons of each lineage’s “original” (0_VH_ and 0_VL_) and “alternative” (A_VH_ and A_VL_) inferred naïve antibodies. In cases where the alternative approach did not infer a naïve sequence that differed from the original naïve sequence identified by partis, alternative chains were paired with original chains, as indicated. For the 157 lineage (C), original and alternative heavy and light chains were also cross-paired. Data are representative of at least two independent experiments. (B) Binding of indicated 157-lineage naïve antibodies (analyte) to monomeric BL035 gp120 (ligand, 250 nM), measured by BLI without the “double reference” approach described in Methods. (D) Graphic summary of paired VH and VL antibody sequences relative to inferred naïve 157-0_VH_0_VK_. Shaded regions demarcate CDRs (gray). Vertical black lines indicate variable region amino acid substitutions. 0: original naïve; A: alternative naïve; m: mature

**Figure 5-figure supplement 1.**
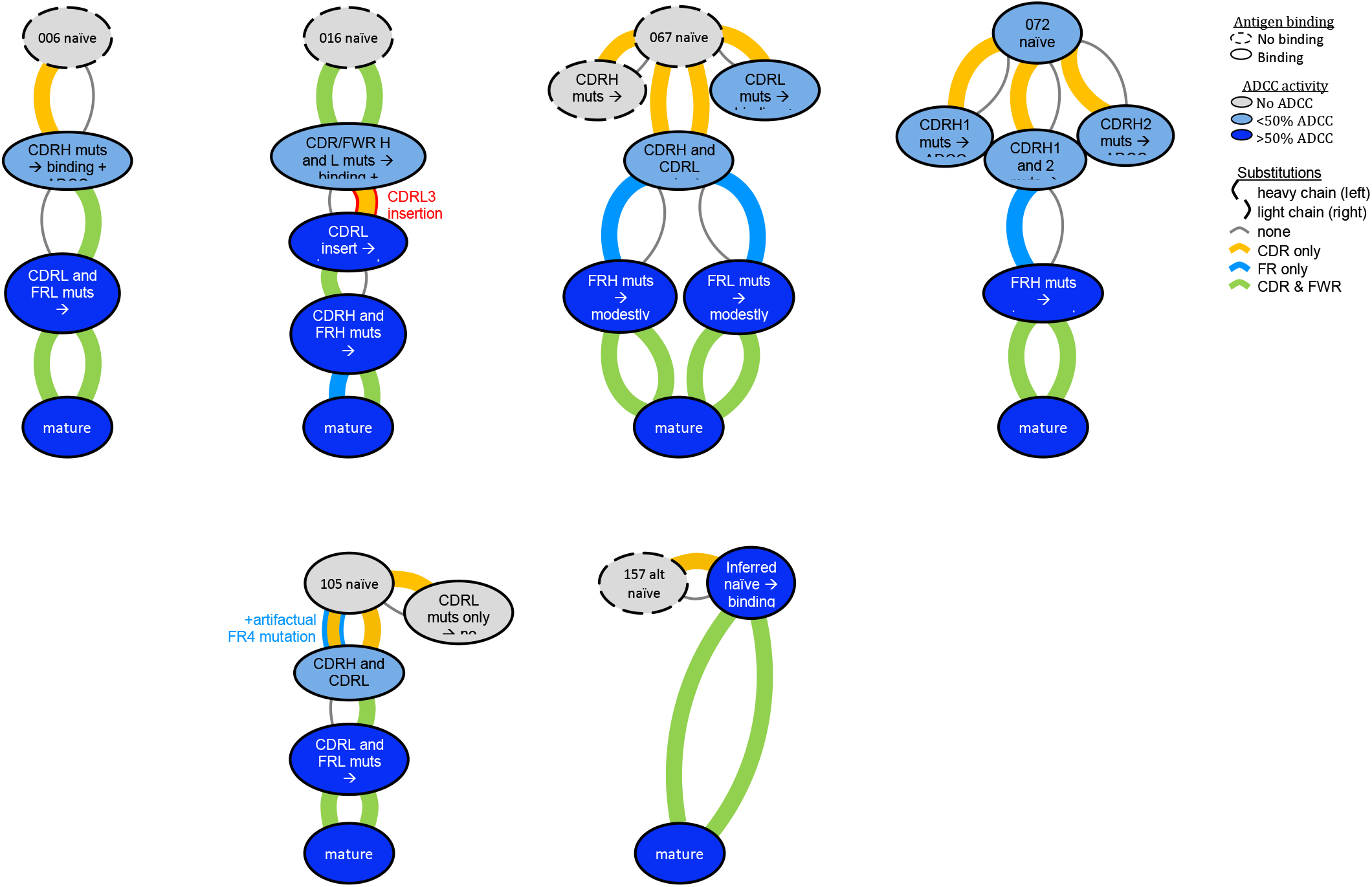
Key steps in ADCC development among six lineages. Graphic summaries of development for each antibody lineage. Antibodies (ovals) are arranged from naïve (top) to mature (bottom) with inferred intermediates in between, colored by ADCC functionality: no function or indeterminate (gray), low function (<50% of mature activity; light blue), high function (>50% of mature activity; dark blue). Arcs connecting ovals indicate amino acid substitutions in the heavy chain (left) and light chain (right), colored by their locality: CDR (yellow), FWR (blue), both (green), or no change (gray, thin). The red outline within the 016 lineage denotes that the light chain mutation is an insertion, not a substitution. The blue outline within the 105 lineage denotes the presence of a FR4 mutation that was likely a computational artifact, since this position backmutates within the mature antibody.

## Acknowledgements

We thank the participants, staff, Scott McClelland, and Ludo Lavreys for continued efforts to oversee the Mombasa cohort. We acknowledge Brian Oliver, Vladimir Vigdorovich, and D. Noah Sather for technical assistance with NGS library preparation and sequence processing. We thank Vrasha Chohan, Haidyn Weight, and Tucker Price for their help with antibody production and testing.

## Competing Interests

The authors declare no competing interests.

## Notes

https://www.ncbi.nlm.nih.gov/bioproject/PRJNA639297/

### Competing Interest Statement

The authors have declared no competing interest.

